# The 3D architecture of the ctenophore aboral organ and the evolution of complex integrative centers in animals

**DOI:** 10.1101/2025.06.26.659588

**Authors:** Anna Ferraioli, Leonid Digel, Daniela Sturm, Jeffrey Colgren, Carine Le Goff, Alexandre Jan, Joan J. Soto-Angel, Benjamin Naumann, Maike Kittelmann, Pawel Burkhardt

## Abstract

The ability to sense and respond to environmental cues is fundamental to animal behavior and survival. In ctenophores - early-branching marine animals - a syncytial nerve net underlies complex behaviors such as geotaxis, feeding, and escape. At the center of this system is the aboral organ (AO), a dense sensory hub that detects motion, light, and pressure and coordinates ciliary movement. However, the AO’s cellular architecture and its integration with the nerve net remain poorly understood. Here, using volume electron microscopy in *Mnemiopsis leidyi*, we reveal that the syncytial nerve net converges and condenses around the AO, forming synaptic connections with diverse effector cells. We annotated 17 distinct cell types, including candidate light and pressure sensors, novel ciliated and secretory cells, and non-synaptic vesicle-rich cells likely involved in volume transmission. Our data shows that signal processing within the AO relies on both synaptic and non-synaptic communication. Gene expression profiling of conserved transcription factors indicates that the AO is a functionally convergent, evolutionarily distinct sensory structure that retains minimal homology. Our findings redefine the ctenophore AO as a highly integrated, multilayered sensory system critical for behavioral regulation.

## INTRODUCTION

Approximately 560 million years ago, the first animals (Metazoa) evolved from unicellular ancestors, giving rise to the astonishing diversity of morphologies and life cycles observed today (1). For over a century, reconstructing the body plan and life cycle of the last common ancestor of the Metazoa (LCAM) has been a central focus in evolutionary biology (2,3). Whether the LCAM possessed an indirect, pelago-benthic life cycle involving a ciliated, diploblastic, free-swimming larva (gastrea) and a benthic adult phase or developed directly, comprising only a single phase, is yet to be determined (3). Well-supported scenarios were challenged by the placement of comb jellies (Ctenophora), rather than sponges (Porifera), as the sister group to all other living metazoan lineages (Fig. 1A) (4–9). This sparked discussions on the independent evolution of some organ systems and cell types from a simple gastraea-like LCAM (3,6,10,11). One of the most controversial topics is the origin of neurons, nervous systems, and their organization together with specialized sensory cells into distinct components of an integrative system (Fig. 1A) (6). Investigating the origin(s) of these systems is crucial to understanding early metazoan evolution and to elucidating whether the LCAM life cycle included a pelagic larva possessing neural structures (3,12,13). Central to this problem is to determine if the aboral sensory organ (AO) of ctenophores is homologous to the pigment ring of some poriferan larvae and the apical organ present in many cnidarian and bilaterian larvae. The majority of extant ctenophores are pelagic, direct-developing invertebrates characterized by a biradial symmetry, tentacles with adhesive colloblasts for prey capture, and a unique swimming mode driven by beating macrocilia organized into eight comb rows (14–16). These comb rows connect to the syncytial subepithelial nerve net (SNN) and, via ciliated grooves, to the AO (11,17,18). The AO develops shortly after completing gastrulation and persists throughout life in all pelagic species (18). It modulates important behaviors such as feeding, vertical migration, and escape responses based on photo-, pressure-, and gravity-sensation (16,19–21). The AO consists of an epithelial floor composed of densely packed multiciliated cells, a statolith suspended by four balancer cilia, and putative secretory, photo-, and pressure-sensitive cells. The entire structure is shielded from the environment by a ring of long ciliated cells forming a dome-like cover (Fig. 1B, 1C). Other aboral sensory organs, located opposite to the blastopore, are found in the larvae of other early-diverging metazoan lineages (22). Some pelagic demosponges possess a pigment ring at their aboral pole that consists of photosensitive long-ciliated cells able to bend laterally that surround a field of mucus-like cells (23–25). Sponge larvae lack neurons, suggesting that cell-cell communication, sensory integration, and behavioral modulation rely on physical contacts and volume transmission of neuroactive molecules (26). Sphere-like swarmer stages described for some placozoans lack any obvious sensory organs (27,28). However, they have only been found in laboratory cultures, and their biological significance within the life cycle is not known. The apical organ of cnidarian planula larvae comprises a diverse assemblage of different sensory neurons, gland cells, and, in some anthozoan larvae, long-ciliated epithelial cells forming an apical tuft (12,29). It is associated with a larval nerve plexus and initiates larval tissue decomposition at the onset of metamorphosis into a benthic polyp (30,31). In *Nematostella vectensis,* the larval apical organ consists of sensory and support cells (32). The apical organ of many bilaterian larvae consists of a conserved set of ciliated sensory neurons, support cells, photo-sensitive cells, and neurosecretory cells (12,32–34). As in cnidarians, it is often associated with a neural plexus (12) and is involved in metamorphosis in the polychaete *Platynereis* (35). The gene regulatory network (GRN), which specifies the anterior-posterior axis and, consequently, the positioning of apical organs, is highly conserved between cnidarians and bilaterians, pointing to the homology of their apical organs (12,36–39). The main actors of the GRN are *Six3/6* and *wnt* genes. Other genes, including *FoxQ, FoxJ, irx, and rx*, specialize in additional specific functions at the aboral end (12,38,39). Overall, the expression of *Six, Fox,* and *rx* genes is located aborally and is essential for the apical organ development (37–39). In parallel, *Wnt* genes form an oral gradient during early stages of development (12,37,38). In contrast, *Wnt* genes in *Mnemiopsis leidyi* are expressed later in development and show an inverse expression pattern confined to specific domains at the aboral pole (40). This suggests that *Wnt* genes are deployed to specify precise aboral domains after the oral-aboral axis is established (40). In addition, *Six* genes in *M. leidyi* appear to have followed an independent radiation (41). Here we used volume electron microscopy to investigate the entire cellular composition and 3D architecture of the AO of the one-day-old ctenophore *M. leidyi.* We uncovered diverse cell morphologies, including previously undescribed ones. We reveal an intimate association with a condensed part of the SNN, forming a potential multilayered circuit combining synaptic conduction and volume transmission. The expression patterns of key genes involved in the GRN of the planulozoan anterior-posterior axis formation suggest a complex evolutionary history of aboral integration centers and provide insights into the life cycle of the LCAM.

**Figure 1.**
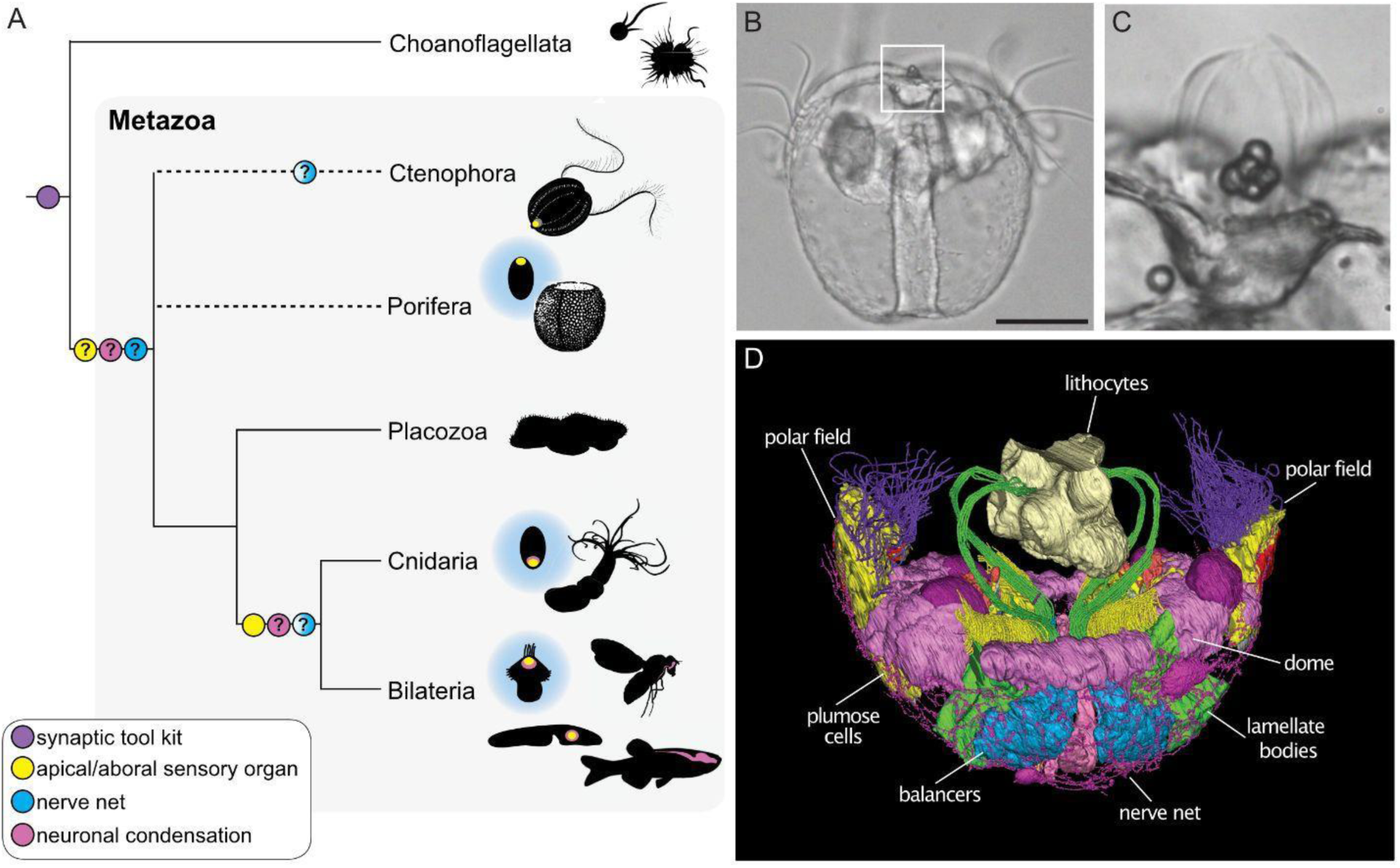
The ctenophore aboral organ and the evolution of nervous system centralization. (A) Metazoan phylogeny and possible scenarios of nervous system origin and centralization. Shaded circles represent alternative scenario (B). Phase contrast image of *M. leidyi* post-hatching cydippid. The white box highlights the aboral organ. (C) Higher magnification of the aboral organ from B, white box. (D) 3D reconstruction of the *M. leidyi* aboral organ main cell types and enclosing nerve net. Scale bar: 50µm.

## RESULTS

### Quantitative 3D morphotype distribution of the ctenophore aboral organ

We provide the first description of the 3D cellular architecture and cell numbers of an early post-hatching ctenophore AO based on the investigation of five serial block face scanning electron microscopy (SBFSEM) datasets (Fig. S1). The segmentation of the nuclei (“M. leidyi RT’’ dataset, Fig. S1) shows that the AO of the one-day-old *M. leidyi* consists of almost 900 cells (Figs. 2A, 2B, 2C, S2) and 17 distinct cell morphotypes (Figs. 2A, S2). These include known cell types such as the balancers, the AO bridge, lamellate bodies, and plumose cells (17,18,20,21,42,43). Novel cell types encompass mono- or multi-ciliated as well as potentially secretory cells.

**Figure 2.**
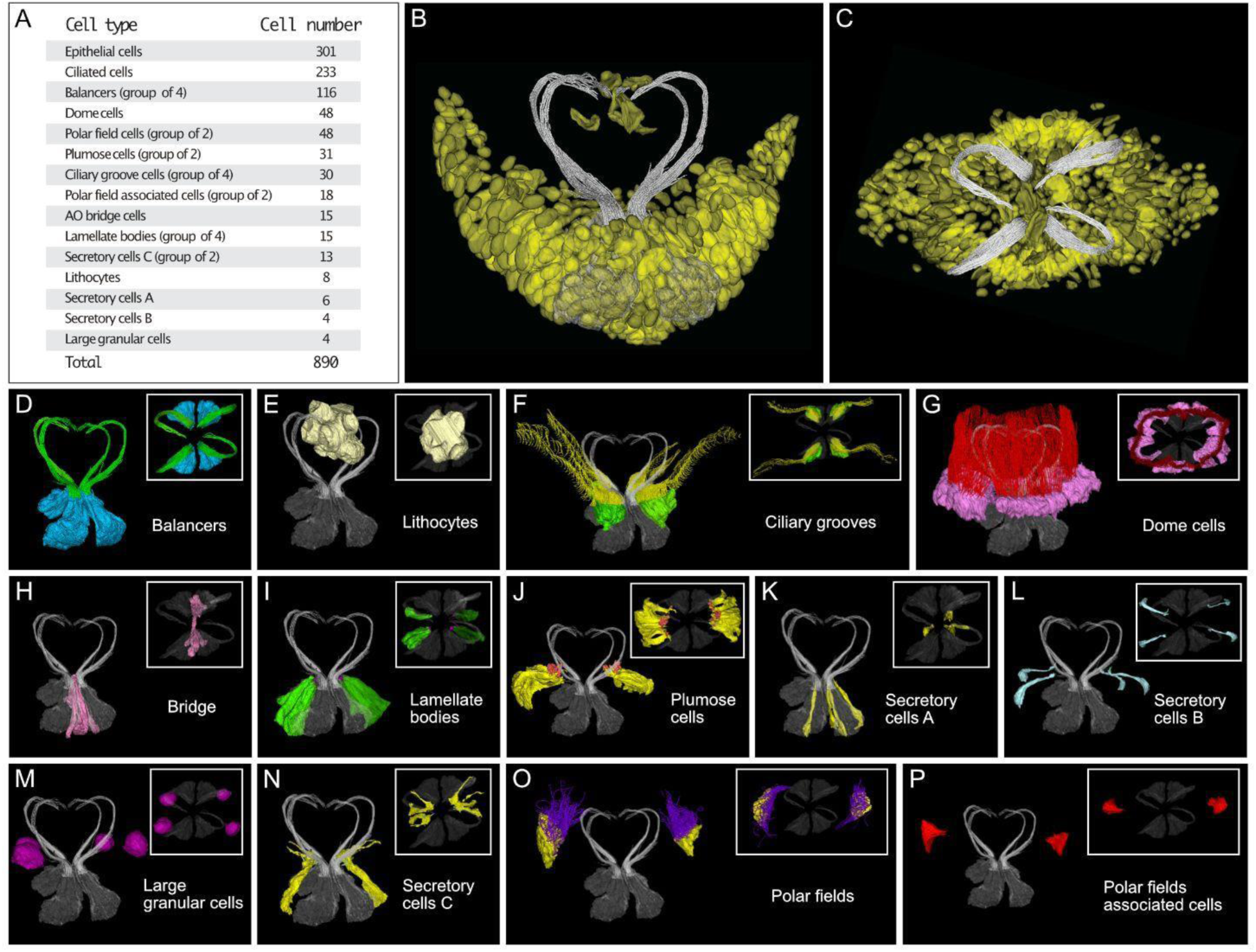
High cell type diversity in the ctenophore aboral organ. (A) Summary of *M. leidyi* aboral organ cell types and cell numbers. (B) Lateral view of the aboral organ nuclei (yellow). (C) Top view of the aboral organ nuclei (yellow). Balancer cilia are gray in (B) and (C). (D) Balancer cell bodies (blue) and sickle-shaped cilia (green). (E) Lithocytes (light yellow) (F). Ciliated groove cell bodies (green) and cilia (yellow). (G) Dome cell bodies (pink) and dome cilia (red). (H) Aboral organ bridge cells (pink). (I) Lamellate bodies (green). (J) Plumose cell bodies (yellow) and cilia (orange). (K) Secretory cells A (yellow). (L) Secretory cells B (light blue). (M) Large granular cells (pink). (N) Secretory cells C (yellow). (O) Polar fields cell bodies (yellow) and cilia (purple). (P) Polar field-associated cells (red). (D-P). Insets: top views. (E-P). Balancers cells and cilia in gray.

The statocyst consists of eight lithocytes suspended on four groups of balancers bearing sickle-shaped cilia (Figs. 2D, 2E, S4 (21,44). The lithocytes feature a large granule and a flattened peripheral nucleus (Figs. 2A, S3B, S2). Each group of balancers is formed by approximately thirty cells (Figs. 2A, S2). Each balancer cell is equipped with a single motile cilium (Fig. S3A (21). The balancers are separated by the AO bridge, consisting of fifteen cells (Figs. 2A, 2H, S2), forming a continuous arch and oriented along the tentacular plane (Fig. S3E (43). The apical sides of each bridge cell are elongated and reach the contralateral group (Fig. S3E). Ciliated grooves originate in proximity to the balancers and are organized into four groups, each consisting of two parallel rows of cells (Fig. 2F). Each group consists of seven to eight multiciliated cells (Figs. 2F, S3C, S2) yielding a total of thirty ciliated groove cells (Figs. 2A, S2). The grooves extend beyond the dome of the AO, bifurcate, and connect to the comb rows (Figs. 2F, S5; (16). The dome consists of forty-eight multi-ciliated cells (Figs. 2A, S2) bearing long, non-motile cilia that protect the AO environment (21), Figs. 2G, S3D). Four groups of one to six cells located in close proximity to the balancers correspond to the lamellate bodies (Figs. 2A, 2I, S2, S3F)(20). Multiple cilia originate basally and extend apically, reaching out to the extracellular space (Fig. S3F). Their cytoplasm harbors numerous small dense-core and electron-lucent vesicles (see below). Morphotypes corresponding to the epithelial papillae or plumose cells (17,19) are arranged into two distal groups from the balancers, and consist of thirteen to eighteen cells (Figs. 2A, 2J, S2). The cell body is sigmoidal, and the cytoplasm exhibits a high density of electron-dense and electron-lucent vesicles (see below). Modified apical cilia show an over-coiled configuration (Fig. S3G).

We discovered multiple putative secretory cells containing distinct types of vesicles. We identified six cells (‘secretory cells A’) neighboring the balancers and harbouring apically polarized electron-lucent vesicles (Figs. 2K, S3H, S2). We reconstructed four cells (‘secretory cells B’) containing small, polarized electron-dense vesicles, and localized in proximity to the pressure cells (Fig. 2A, 2L). The cell bodies are slender, the apical and basal poles are enlarged, and the nuclei are located basally (Fig. S3I). We annotated thirteen cells (‘secretory cells C’) distributed in two groups in proximity to the balancers, bearing long apical protrusions and small polarized electron-lucent vesicles. (Figs. 2A, 2N, S3K, S2). Four large granular cells are located at the edge of the AO, close to the dome cells (Figs. 2A, 2M, S2). The cell body is rounded, the nucleus is located basally, and several types of vesicles and reflective inclusions are densely interspersed within the cytoplasm (Fig. S3J). Most of the remaining AO cells bear one or multiple cilia (Table S1) and form a symmetrical pattern (Fig. S6A). We identified about two hundred mono-, bi-, and tri-ciliated cells (Figs. 2A, S2, Fig. S3L to N). These consist of elongated cell bodies, few mitochondria, and large nuclei (Fig. S3L to N). The large majority of these cells are mono-ciliated cells (Fig. S6A, B), whereas bi- and tri-ciliated cells are distributed in patches (Fig. S6B, S6C). Among the mono-ciliated cells, we identified two single cells bearing longer cilia, situated among the ciliated grooves on the tentacular plane (Fig. S3O). Live tubulin staining of cydippid indicates that these are motile cilia and bend laterally toward the ciliated grooves, establishing apparent contact (Fig. S6D, S6E, Videos S1, S2). A cluster of monociliated cells (‘bundle cells’) is localized centrally, underneath the bridge (Fig. S3P, S6F). We annotated forty-eight polar field cells (Figs. 2A, 2O, S2) and eighteen polar field-associated cells (Figs. 2A, 2P, S2) distributed in two groups on the outer side of the dome (Fig. 1D). The polar field cells are multiciliated cells equipped with long motile cilia (Fig. S2Q), whereas the polar field-associated cells are non-ciliated, displaying thin apical protrusions (Fig. S2R). Finally, we annotated about three hundred round-shaped and elongated epithelial cells (Figs. 2A, S2, Fig. S3S, S3T) scattered amongst other AO cells (Table S1).

### Condensation of the subepithelial nerve net around the aboral organ

We traced neural cell bodies and neurites of the SNN near the AO and consistently observed a syncytial architecture with interconnected neurites extending from the neural cell bodies as previously described (11)(Fig. 3A). The AO-adjacent nerve net consists of four symmetrically distributed neural cell bodies (Fig. 3A). The nerve net encases the AO, and the neurites are situated close to the basal poles of the cells (Fig. 3A, Fig. 3A-top inset, Fig. 3A-bottom inset). The morphology of the neurites shows the typical pearl-on-a-string structure (Fig. 3A-bottom inset) and is overall consistent with that of the subepithelial nerve net (11)(Fig. 3B, 3C). However, we observed some neurites exhibiting slightly thicker and continuous morphologies, interspersed with the pearl-on-a-string structures (Fig. 3C). Similar to the subepithelial nerve net, the neural cell bodies and neurites contain dense-core and electron-lucent vesicles (Figs. 3B, 3C, S3U, S7). Interestingly, we observed sheets of endoplasmic reticulum (ER) embedded in the cytoplasm of these cell bodies, suggesting a higher synthetic activity in this cellular compartment. This is consistent across datasets (Figs. 3, S7). The datasets of “animal 4” and “animal 5” (Fig. S1) revealed neural cell bodies containing two nuclei (Fig. 3D, 3E, insets), although the ultrastructure is similar to mono-nucleated cell bodies, including vesicles and ER membranes (3D, 3E). Extended reconstruction of the SNN towards the epithelium flanking the AO revealed a change in the SNN architecture. While the syncytium is maintained, we observed a substantial increase in neurite density in close proximity to the AO cells. Here, neurites are densely packed, highly interconnected, and the number of polygons per SNN area increases. We define this configuration as “condensed” SNN. This is characterized by the presence of denser polygons in contrast to the SNN. (Fig. 3F, 3G). The “condensed” and “regular” SNN configurations are observed consistently across datasets (Fig. 3F, 3G).

**Figure 3.**
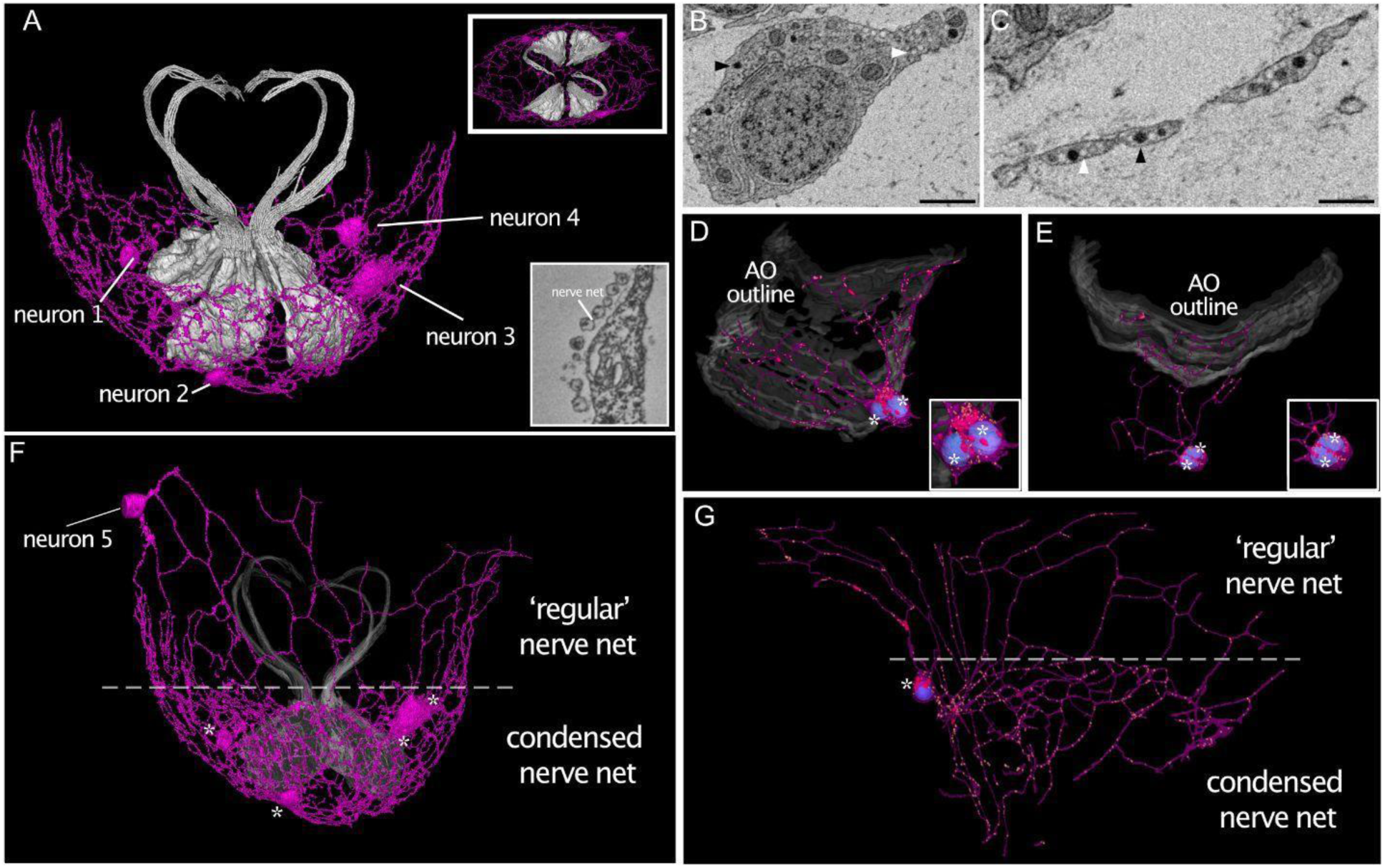
Condensation of the nerve net encasing the aboral organ. (A) 3D reconstruction reveals the syncytial architecture of the aboral organ nerve net (pink). Four nerve net cell bodies are interconnected by neurites. The relative position to the balancer cells (gray) is shown (top inset: top view of the balancer cells and the aboral organ nerve net. Bottom inset: ultrastructure of the “pearls-on-a-string” morphology. (B) SBFSEM cross section of a nerve net cell body, including electron-dense (black arrowhead) and electron-lucent vesicles (white arrowhead). (C) SBFSEM cross section of a neurite including electron-dense (black arrowhead) and electron-lucent vesicles (white arrowhead). (D and E) 3D reconstruction of nerve net cell bodies containing two nuclei (white asterisks). The neurites extend toward the aboral organ cells (AO outline, in grey). (F) The nerve net above the dashed line extends towards the neighboring epithelial cells and shows characteristic, large polygons, including an additional cell body (neuron 5). The nerve net below the dashed line, which includes four cell bodies (asterisks), shows compact, small polygons. (G) 3D reconstruction of high-pressure frozen tissue confirms condensation of the nerve net encasing the aboral organ. Nerve net cell body (asterisk: nerve net cell body, blue: nucleus and neurites extending from it (pink). Yellow: electron-dense vesicles, red: mitochondria. Scale bars: 1µm.

### Signal conduction routes in the aboral organ

Our 3D reconstructions revealed neurite projections that extend from SNN neurons to form synaptic contacts onto two effector cells, the balancers and the bridge cells (Fig. 4). Multiple synapses were detected at the base of the balancer cells, consistent with previous observations (45) while some neurites extended amongst the balancer cells and reached towards the apical poles of the cell complex (Fig. 4A and B). The neurites extend from a proximal neural cell body and envelop the group of balancers (Fig. 4C, 4D). A similar multi-synaptic configuration is detected at the bridge cells, although the neurites are not intertwined within the cell group (Fig. 4E, 4F). The synaptic contacts are characterized by the typical synaptic triad, including a mitochondrion, ER, and vesicles (Fig. 4G) (42,45). Distinct and specialized morphotypes include the lamellate bodies, plumose cells, and putative secretory cells (Fig. 4H to 4L). These are equipped with numerous non-synaptic electron-dense and electron-lucent vesicles (Fig. 4H to K). The lamellate bodies harbour small electron-lucent vesicles spanning the length of the cell body and medially polarized small electron-dense vesicles (Fig. 4H). The plumose cells’ cytoplasm is heavily loaded with large electron-dense vesicles, whereas electron-lucent vesicles polarize at the apical pole (Fig. 4I). In contrast, the secretory cells contain only one type of non-synaptic vesicle (Fig. 4J to 4L). Secretory cells A and C contain non-synaptic electron-lucent vesicles (Fig. 4J, 4L). In both cases, the vesicles are polarized apically. Conversely, secretory cells B harbour only electron-dense vesicles, which show a medial-basal polarization (Fig. 4K). These cell types may contribute to an alternative non-synaptic route for signal conduction in the AO through volume transmission (Fig. 4M).

**Figure 4.**
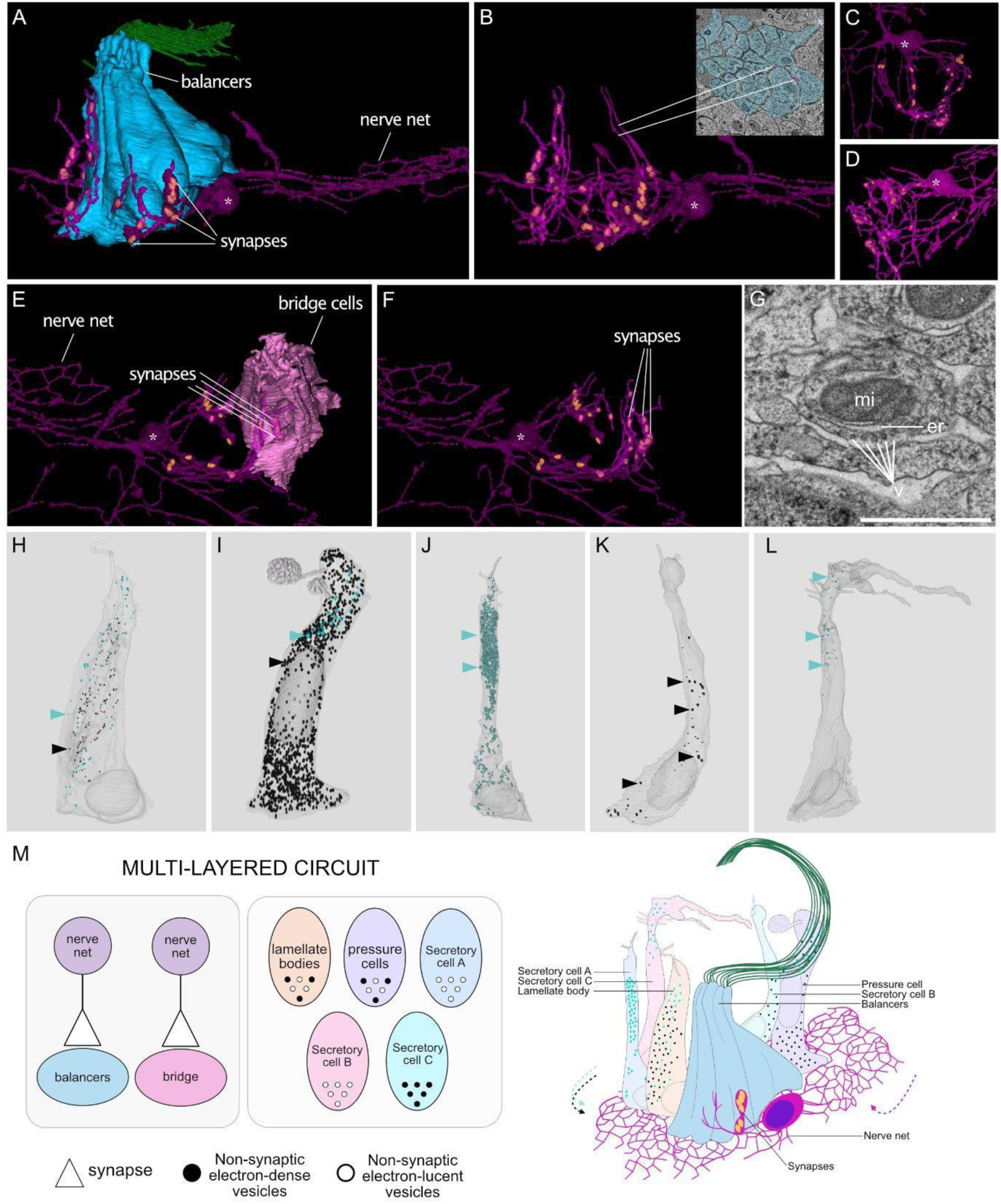
Multi-layered neuronal architecture employed by the ctenophore aboral organ. (A) 3D reconstruction reveals direct synaptic contacts between nerve net neurons and balancers. Blue: balancers, green: cilia. (B) Long neurites containing multiple synapses are intertwined with the balancers’ cell bodies. Inset: SBFSEM cross-section. Blue balancers, pink: nerve net neurites. (C) Top view and (D) lateral view of the nerve net associated with balancers. (E) 3D reconstruction reveals direct synaptic contacts between nerve net neurons and bridge cells. (F) Several synapses form connections to bridge the cell bodies. (A-F) Asterisks: nerve net cell bodies, yellow: mitochondria. (G) TEM cross-section of a synaptic triad, including a mitochondrion (mi), endoplasmic reticulum (er), and synaptic vesicles (v). (H) 3D reconstruction of a lamellate body. (I) 3D reconstruction of plumose cells (J) 3D reconstruction of secretory cell A (K) 3d reconstruction of secretory cell B (L) 3D reconstruction of secretory cell C. Light blue dots and arrowheads: non-synaptic electron-lucent vesicles; black dots and arrowheads: electron-dense vesicles. (M) Aboral organ circuitry and circuit model indicating synaptic contacts and potential volume transmission. Scale bars: 1µm.

### Anterior-posterior patterning genes in *M. leidyi*

We investigated the expression of genes involved in the conserved planulozoan anterior-posterior GRN. Specifically, we searched for orthologs of the *Six3/6* family in *M. leidyi*. Using orthology inference analysis and gene phylogeny, we identified two genes, *ML01466a* and *ML082610a,* sharing a common ancestor with the *Six3/6* clade (Table S2, Fig. S9). These genes (‘*MlSix3/6A’* and ‘*MlSix3/6B’,* Fig. S8) show a close relationship, and their grouping in the phylogenetic tree suggests a possible lineage-specific evolutionary event, consistent with previous studies (41)(Fig. S9). In the one-day-old *M. leidyi, MlSix3/6A* is expressed along the comb rows, and *MlSix3/6B* is expressed in the pharynx area and large, isolated cells on the outer epithelium of the tentacle bulb and the distal pharynx area, or mouth (Fig. S8B). The temporal expression pattern shows that both genes start to be expressed around 8 - 11 hours post-fertilization (Fig. S8A)(46). We addressed the homology of the ciliary structures of the AO with those of planulozoan ciliary tufts, focusing our gene orthology analysis on the Forkhead transcription factor, *FoxJ.* Our analysis revealed orthology of *ML23712a* (‘*MlFoxJ*’) with other genes belonging to the *FoxJ* family (Table S2, Fig. S10). Interestingly, in the one-day-old *M. leidyi, MlFoxJ* is expressed in the AO in four dense symmetrical patches, and a punctate signal can be detected in between the dense areas and in the ciliated grooves area connecting to the comb rows as well (Fig. 5A, 5B, 5C, 5E). A well-defined expression of *MlFoxJ* is detected along the comb rows (Fig. 5B, 5D, 5E) that develop around nine hours after fertilization (47). Temporal expression data indicate that *MlFoxJ* starts to be expressed around gastrulation.

**Figure 5.**
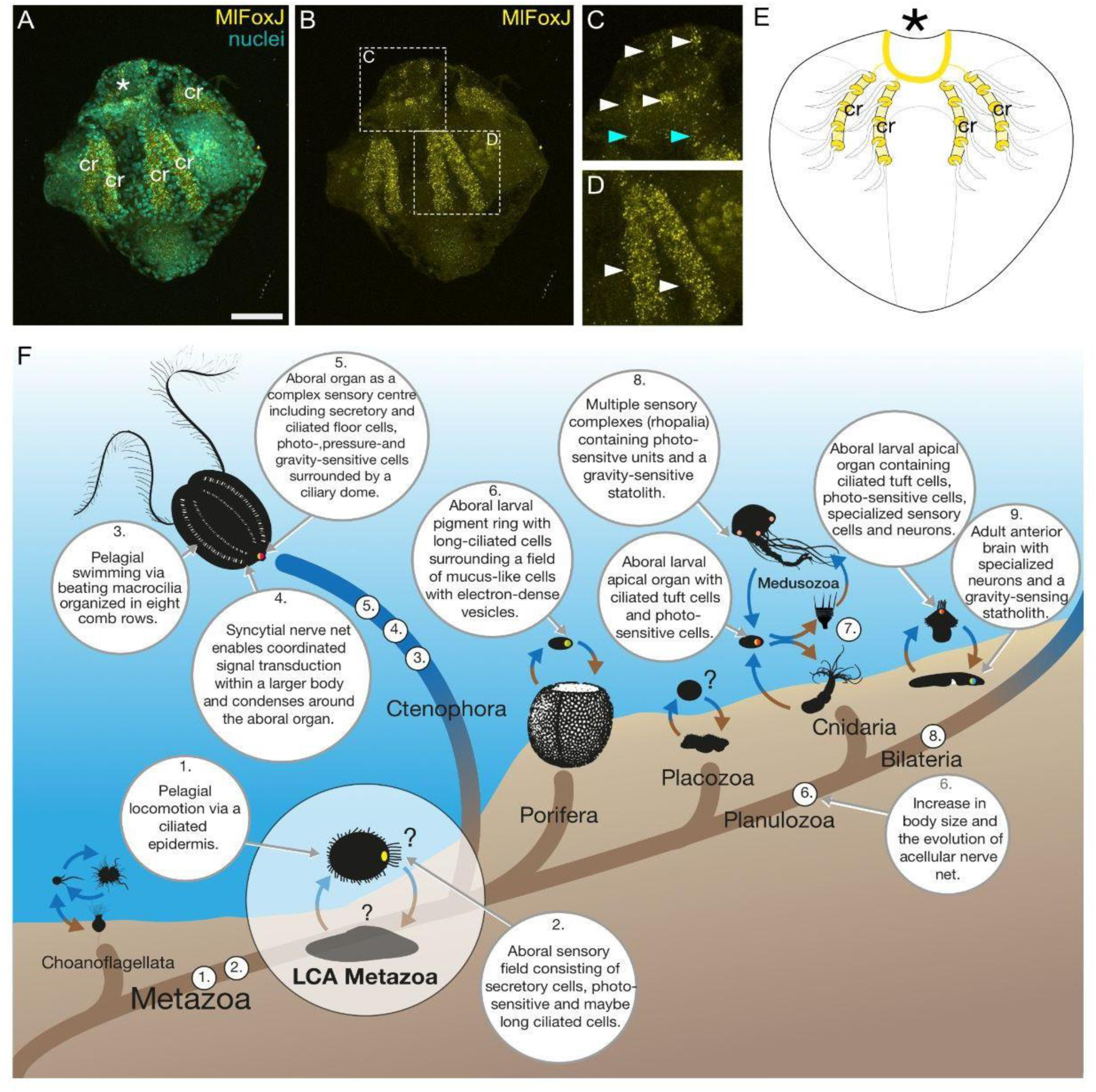
Complex evolutionary history of aboral integration centers. (A) Expression of *MlFoxJ* (yellow) and nuclei (cyan) in a one-day-old cydippid of *M. leidyi.* (B) Expression of *MlFoxJ* in the aboral organ and the comb rows. (C) Higher magnification showing the expression of *MLFoxJ* in the aboral organ. White arrowheads indicate the brightest areas of expression in the AO. Cyan arrowheads indicate expression in the ciliated grooves. (D) Higher magnification showing the expression of *MlFoxJ* in the comb rows. White arrowheads indicate the comb rows. (E) Schematics of the expression of *MlFoxJ* in the one-day-old cydippid of *M. leidyi.* Asterisks indicate the position of the AO, cr: comb rows. Scale bar: 50 μm. (F) Schematics explaining a potential evolutionary scenario of sensory complexes in the last common ancestor of Metazoa (LCAM).

## DISCUSSION

### Aboral organ complexity in ctenophores

The assemblage of integrated sensory complexes in animals likely marked the expansion of the pelagic phase within animals. The enhanced capability of interpreting the environment on a large scale allowed early animals to process inputs and react rapidly (48,49). Since its first description as a sensory organ by Richard Hertwig in 1880, several studies have aimed to understand the unique morphology and behavioral modulation of the ctenophore AO (16,18,42–44). Our volume electron microscopy-based 3D reconstruction of a one-day-old *M. leidyi* reveals the spatial organization of the cellular diversity within the AO. We show that the AO consists of about 900 cells, distributed into 17 morphotypes (including previously described balancers, bridge cells, plumose cells, and lamellate bodies). In addition, our study describes putative secretory cells, ciliated cells, epithelial cells, dome, polar fields, and polar fields-associated cells, and a previously undescribed association with a condensed part of the SNN. Moreover, we report novel morphotypes of several secretory and ciliated cells. Distinct vesicle populations in secretory cells indicate discrete signals and signalling pathways (see below), whereas we hypothesize that so far undescribed ciliated cells contribute to the overall sensory capability of the AO. Altogether, the presence of multiple and distinctive morphotypes identifies the AO as a complex sensory organ capable of processing and integrating multimodal sensory signals. A direct connection with the locomotory system (i.e., the comb rows) establishes the AO as an essential system for input processing and behavioral coordination.

### Condensed neurons are associated with the aboral organ complex

The AO complex is enclosed by a condensed section of the SNN (Fig. 3). Interestingly, the cytoplasm of the SNN cell bodies is enriched with ER sheets compared to previous findings (11)(Fig. S7), indicating high synthetic activity. Additionally, we found SNN cell bodies with multiple nuclei in slightly older animals (Fig. 3), which could result from karyokinesis, adding insights into how the neural cell body number expands and the SNN remodels in older animals. The condensed architecture points towards the necessity to enhance neuronal wiring in an area of increased signalling activity. Alternatively, though not in contrast, a hierarchical mode of signalling mechanisms could exist, whereby the input, processing, and output take place in the AO compartment and involve the concerted participation of specialized cells and the SNN. The output likely propagates to the rest of the body, yielding behavioral modulation (e.g., change of the swimming pattern in relation to light intensities (16,50)).

### An emerging multilayered circuit

We report synaptic connections from the SNN onto the balancers and the bridge. Previously reported synaptic contacts from the SNN onto the ciliated grooves (51) and comb cells (11) support the possibility of a joint neural circuit that coordinates swimming responses (16,21,52). Alternatively, nerve net synapses contacting the bridge cells could form a different sub-circuit involved in prey-capture behaviours (43). Additional synaptic contacts onto lamellate bodies and secretory cells have previously been reported for adult ctenophores (19). These contacts were not present in our data, indicating that lamellate bodies, and possibly other cell types, become wired into one or multiple circuits later in development, reflecting an even greater behavioral coordination. In any case, synaptic conduction constitutes a defined pathway for signal propagation towards the AO. However, the propagation via volume transmission appears to be functioning simultaneously. Lamellate bodies, plumose cells, and other secretory cells harbour highly polarized, electron-dense and electron-lucent vesicles (Fig. 4) and could serve as key players in volume transmission (Fig. 4L). The occurrence of volume transmission in the AO is supported by the expression of neuropeptides and their effect on swimming behaviors (e.g. ML065755) (51). The coexistence of synaptic and volume transmission suggests that ctenophores utilize a multilayered circuit to connect the nervous system and cellular components in the AO (Fig.4L). To further dissect neural signaling in ctenophores, a better understanding of the molecular signature of the chemical synapses and a map of neuropeptide-receptor interactions is required.

### Evolutionary origin of sensory complexes

Comparative molecular and morphological studies established the homology of cnidarian and bilaterian apical organs (12,33,36). Whether the ctenophore AO is homologous to these remains controversial, due to its enhanced morphological complexity (fully illustrated in this study) and lack of consistency of expression of conserved genes and transcription factors (40,48). We further addressed the homology of *M. leidyi* AO with that of planulozoan by exploring gene expression patterns for the orthologs of the highly conserved transcription factors *Six3/6, FoxQ2,* and *FoxJ1* in 1-day-old *M. leidyi.* We could not detect expression of orthologs of *MlFoxQ2* in 1-day-old *M. leidyi,* and along with previously reported differences in expression data for *Wnt* family genes (40), our data indicate that the classic AP-GRN is not conserved (Figs. 5A, S8). This suggests that the evolution of this GRN occurred sometime after the branching of ctenophores from the planulozoan ancestor. The emergence of *Wnt*-inhibitor expression at the aboral pole in the planulozoan ancestor may have enabled an expanded GRN and the development of a new apical neuroectodermal plate (12,36), including the specification of neurons around the ancestral apical organ. In ctenophores, persistent aboral *Wnt* expression may have prevented the formation of a similar neuroectodermal plate and therefore neurons and chemical synapses, favoring instead volume transmission as the primary signaling mode in the apical organ. Notably, ctenophore comb rows develop in a *Wnt*-free zone and share GRN components with the planulozoan apical organ and plate, including *FoxJ*, *Six3/6A* (this study), and *SoxB1* (53), suggesting a possible convergent evolution of similar GRNs for comb row and apical plate development. *FoxJ1* transcription factor orthologs are expressed in the apical organ of *M. leidyi*, early pigment-ring forming embryos of *A. queenslandica* (54), and the apical region of *N. vectensis* (38) and *P. dumerilii* larvae (12). The occurrence of *FoxJ1* predates the evolution of animals. It is downregulated in cRFXa mutants with aberrant ciliogenesis, although direct perturbation of *FoxJ1* has no effect on ciliogenesis in the choanoflagellate *S. rosetta* (Coyle et al, 2023). This indicates its ancestral involvement in ciliated structures, but also its tight regulation by other factors. The expression of *MlFoxJ* in both the AO and the comb rows in *M. leidyi* suggests that the involvement in ciliary structures is conserved. However, we can not determine whether *MlFoxJ* plays a role in the patterning of ciliated structures or belongs to a terminal selector module that has been redeployed in the comb rows as well as the AO of ctenophores. We propose that a specialized apical sensory spot evolved in the LCAM, possibly involving the expression of *FoxJ1* and later diversifying across early/deep-branching metazoans (Fig. 5F). We hypothesize that the LCAM life cycle included a pelagic phase with an apical sensory spot likely composed of cilia, photo-receptive, and secretory cells as proposed by Marlow et al, (12) for the last common ancestor of planulozoan (Fig. 5F). It became a coordination center in ctenophores, larval poriferans, and planulozoans by associating with different functions (e.g., gravity, light, pressure sensation, metamorphosis). In ctenophores and the planulozoan ancestor, larger epithelialized bodies and more active lifestyles placed similar selective pressure for better sensory integration. This may have led to the independent evolution of a larval neural plexus and adult nerve net in planulozoans (12), and to a condensed syncytial SNN in ctenophores (this study; Jokura et al., 2025). In bilaterian ancestors, the integrated apical organ may have served as a nucleation center for an adult brain, which relocated to the oral pole and became a new command center (12,48). Our interpretation, that a pelagic phase with an apical ciliated spot could be homologous among ctenophores, poriferans, cnidarians, and bilaterians, challenges the successive addition of larval stages (larval intercalation hypothesis) and the idea of a direct-developing LCAM (e.g., (55). Increased taxon sampling of early-branching animals and further investigation of ciliary GRNs across species will clarify how the patterning of ciliated sensory complexes evolved. Our study provides a fully resolved ultrastructural cell type atlas of the AO of a 1-day-old *M. leidyi*. It sheds light on the neural circuitry underlying ctenophore behavior and highlights the relationship between cell type diversity, multimodal circuitry, and neural condensation in one of the earliest evolved animal sensory systems.

## MATERIALS AND METHODS

### Animal husbandry

For electron microscopy, *M. leidyi* cydippids were maintained in 300 mL glass beakers and fed daily with *Brachionus,* as previously described (56) (68). For Hybridization Chain Reaction (HCR), adults of *M. leidyi* were maintained in 25 L kreisels and fed daily with *Brachionus* and *Artemia.* In both cases, animals were maintained under a 17h/7h light-to-dark cycle (56) (68). Following 7 hours of darkness, adults were transferred to 300 ml beakers and allowed to release gametes. Fertilized eggs were transferred to 1 L glass beakers with aeration and incubated until fixation.

### Electron Microscopy sample preparation

Fixation by high-pressure freezing (HPF) and freeze substitution (FS) was previously described (51). Briefly, *M. leidyi* cydippids were frozen in 20% BSA in natural seawater in 0.2 mm aluminum planchettes in a Baltech10 High Pressure Freezer. Samples were placed in cryotubes with frozen 0.1% UA + 1% osmium tetroxide in liquid nitrogen and transferred to an RMS freeze substitution machine: 20 hr at −90°C, 21 hrs from −90°C to −20°C, 24 hr at −20°C, 5 hrs from −20°C to 4°C, 30 min from 4°C to 20°C. At room temperature samples were washed in acetone and incubated in 1% tannic acid in acetone for 2 hr, washed in acetone, incubated for 2 hrs in 1% osmium tetroxide, washed in acetone and infiltrated in a series of 10%, 30%, 50%, 70% Taab 812 hard Epoxy resin for at least 3 hour each or overnight, then infiltrated 3x in fresh 100% Taab 812 hard Epoxy resin for at least 4 hrs each or overnight at 4°C. Cydippids were then embedded in fresh resin and polymerized at 70°C for at least 24 hrs.

For chemical fixation, *M. leidyi* cydippids were fixed in 2% glutaraldehyde and 2.5% paraformaldehyde in 0.1 M NaCac pH 7, made up in 27 ppt seawater, gently rocking for 3 hrs at 4°C. Cydippids were then washed 3x in seawater, then stained with 2% Osmium (aq.) overnight at 4°C, washed 3x in water, stained with 1% Tannic Acid (aq.) for 4 hrs at 25°C washed 3x in water, stained with 1% Osmium overnight at 4°C, washed 3x in water, stained with 2% Uranyl acetate (aq.) for 2 hrs, washed 3x in water and dehydrated in a series of 10%, 30% and 50% EtOH for 45 minutes each. After overnight 70% EtOH incubation, they were placed in 4x fresh 100% Ethanol for 4 hours or overnight each, and finally infiltrated in a series of 10%, 30%, 50%, 70% for at least 3 hours each, then infiltrated with 812 Epoxy resin as described above.

### SBFSEM

For Serial Block Face SEM, *M. leidyi* cydippids were prepared as previously described (51). Briefly, samples were glued onto SBFSEM stubs with conductive Epoxy resin (Chemtronics, Hoofddorp, Netherlands), trimmed with a diamond trimming knife, and sputter-coated with 20 nm gold. Images were collected with a Merlin Compact SEM (Zeiss, Cambridge, UK) with the Gatan 3View system and Gatan OnPoint BSD and Focal Charge Compensation (100%) with a 20 μm aperture, at 1.8 kV and 1 μs pixel time. Pixel size of raw data is specified in Fig. S1 for each dataset. The datasets are available here: 10.6084/m9.figshare.29314115.

### SBFSEM datasets overview, segmentation and 3D reconstruction

We generated 5 serial block face scanning electron microscopy (SBF-SEM) datasets using chemical fixation (“M. leidyi RT”, Fig. S1) and high-pressure freezing (“M. leidyi HPF1”, “M. leidyi HPF2”, “M. leidyi HPF3” and “M. leidyi HPF4”, Fig. S1). SBF-SEM sections were imported into Fiji (57) and aligned to generate z-stacks using default parameters, as previously described (11,51). Stacks were imported into TrakEM2 (58), and segmentation of organelles, cilia, and nerve net was performed manually. Segmentation of nuclei and membranes was performed semi-automatically by using the function ’interpolate gaps’, where possible. Meshes were visualized with 3D Viewer (59), rendered at a resolution of 1 to 3 for the nerve net, of 5 for single cells, and 10 for composites. Meshes were smoothed by a factor of 5 to 10, and small vesicles were not smoothed. We exploited ‘animal 1’ dataset the most, as it contains a full coverage of the AO. We leveraged the remaining datasets to examine details of the subcellular features and the nerve net architecture.

### Fixation of specimens for whole mount *in situ* hybridization (HCR)

Cydippids of *M. leidyi* were collected after 24 hours post-fertilization and fixed as previously described by Mitchell et al. (2021) with the following modifications. Briefly, animals were transferred into glass vials containing 2.5mL of artificial seawater (ASW) and rapidly relaxed by adding 500uL of 1M MgCl2. Fixation was conducted with 16% ice-cold Rain-X® in artificial sea water (ASW, 27ppt, pH=8.2) for 20 min under constant rotation. Cydippids were post-fixed in 3.7% ice-cold Formaldehyde solution in ASW for 20 minutes at 4°C under constant gentle rotation. Following fixation, animals were washed 3 times in Phosphate saline buffer (PBS, 137 mM NaCl, 2.68 mM KCl, 10.14 mM Na2HPO4, 1.76 mM KH2PO4, pH 7.4) with 0.1% Tween 20 (Sigma Aldrich) and dehydrated stepwise in methanol. Fixed specimens were stored at -20°C and rehydrated stepwise in PBS-0.1%Tween 20 before proceeding with HCR.

### Whole mount *in situ* hybridization (HCR)

In situ HCR v3.0 with split initiator probes was performed using probe sets, amplifiers, and buffers obtained from Molecular Instruments (https://www.molecularinstruments.com) (61). Probes for HCR were designed with the insitu_probe_generator custom script (62). Probe sets were ordered from Integrated DNA Technologies (https://eu.idtdna.com). Rehydrated samples were incubated in the Probe Hybridization buffer (Molecular Instruments) for 1 hour at 37°C. Probes were used at a concentration of 16nM and incubated overnight at 37°C. Following incubation, samples were washed in Probe Wash buffer (Molecular Instruments) 4 times for 15 minutes at 37°C, then 5 times in 5x saline sodium citrate buffer (SSC, 0.75M Sodium Chloride and 0.075M Sodium Citrate) containing 0.1% Tween 20 (Sigma Aldrich) for 5 minutes at room temperature (RT). Samples were pre-amplified in Amplification buffer for 30 minutes at RT. Hairpins were used at a concentration of 60nM, heated in a PCR thermocycler for 90 seconds at 95°C, and allowed to cool down for 30 minutes at RT in the dark. Hairpins were then mixed with the Amplification buffer, and solutions were added to the samples and incubated overnight at RT in the dark. Following amplification, samples were washed twice in 5x SSCT for 5 minutes, followed by nuclear staining using 1:2000 Hoechst 33342 (10mg/ml, ThermoFisher Scientific) for 30 minutes at RT. Finally, samples were washed 3 times in 5x SSC and mounted in SlowFade Glass™ soft-set antifade mounting medium.

### Live staining

Recently hatched cydippids were washed with ASW and incubated with 1uM SiR tubulin (Cytoskeleton, Inc.; CY-SC002) in ASW overnight at 18°C. The animals were then washed 3 times with ASW before mounting on glass bottom dishes (MatTek; P35G-1.5-10-C) in 0.5% low-melt agarose (Cat#CR-6351.5) in ASW containing 100nM SiR Tubulin.

### Confocal imaging and image analysis

Images of fixed specimens and videos of live animals were acquired with an Olympus FV3000 confocal laser scanning microscope, and Z-stacks maximum intensity projections were obtained with Fiji (57). Contrast and brightness of the images were adjusted by refining the contrast and the brightness of the maximum projections to improve the visualization of the gene expression.

### Gene orthology

Search for clusters of orthologous genes was performed with Broccoli 1.2 (63) using the Maximum Likelihood algorithm for steps 1, 2, and 3. We used available proteomes for *Homo sapiens* (UP000005640_9606), *Drosophila melanogaster* (UP000000803_7227), *Caenorhabditis elegans* (UP000001940_6239), *Nematostella vectensis* (UP000001593_45351), *Mus musculus* (UP000000589_10090), *Danio rerio* (UP000000437_7955), *Platynereis dumerilii* (901 annotated proteins) and *Mnemiopsis leidyi* (https://research.nhgri.nih.gov/mnemiopsis/). We recovered two orthogroups, each containing *MlFoxJ and MlSix3/6A and B* (Table S2). To validate our orthogroups, we performed gene phylogenies. We compiled lists of protein families from the sample of species used in Broccoli, including only full-length proteins. We generated sequence alignments using MUSCLE 5.3 (64) with default parameters. Alignments were cleaned using TrimAl 1.5 (65) with default parameters. Gene phylogenetic trees were built using IQTree2 2.4 (66) using the MFP option, allowing for the best substitution model to be selected. Phylogenetic trees including *MlFoxJ and MlSix3/6A, MlSix3/6B* are available in Figures S9, S10.

## ACKNOWLEDGEMENTS

High-pressure freezing and SBFSEM, and TEM imaging were conducted in the Oxford Brookes Centre for Bioimaging. This work was supported by the Michael Sars Centre core budget, the European Research Council Consolidator Grant (101044989, “ORIGINEURO”), and the Human Frontier Science Programme (RGP025/2023). We thank Roberto Feuda for discussion on apical gene regulatory networks in animals, Stefan Richter and Christian Wirkner for discussions on metazoan larvae, and Burkhardt lab members for discussion and feedback on the study.

## Notes

### Competing Interest Statement

The authors have declared no competing interest.

