## Supplementary Information for "The 3D architecture of the ctenophore aboral organ and the evolution of complex integrative centers in animals"

| | animal 1 | animal 2 | animal 3 | animal 4 | animal 5 |
|---|---|---|---|---|---|
| fixation | chemical | HPF | HPF | HPF | HPF |
| resolution | 5 x 5 x 100 nm | 3 x 3 x 100 nm | 5 x 5 x 100 nm | 5 x 5 x 100 nm | 5 x 5 x 100 nm |
| name | M. leidyi RT | M. leidyi HPF 1 | M. leidyi HPF 2 | M. leidyi HPF 3 | M. leidyi HPF 4 |
| coverage | complete | partial | partial | partial | partial |

**Figure S1.** SBFSEM datasets used in this study

| Cell type | Cell number | |
|---|---|---|
| ***1. Ciliated cells*** | | **207** |
| mono-ciliated cells | 142 | |
| bi-ciliated cells | 61 | |
| tri-ciliated cells | 4 | |
| ***2. Epithelial cells*** | | **155** |
| ***3. Epithelial elongated cells*** | | **146** |
| ***4. Balancers*** | | **116** |
| group 1 | 30 | |
| group 2 | 27 | |
| group 3 | 28 | |
| group 4 | 31 | |
| ***5. Dome cells*** | | **48** |
| ***6. Polar fields cells*** | | **48** |
| field 1 | 20 | |
| field 2 | 28 | |
| ***7. Plumose cells*** | | **31** |
| group 1 | 18 | |
| group 2 | 13 | |
| ***8. Ciliated groove cells*** | | **30** |
| group 1 | 8 | |
| group 2 | 8 | |
| group 3 | 7 | |
| group 4 | 7 | |
| ***9. Bundle cells*** | | **26** |
| ***10. Polar fields associated cells*** | | **18** |
| group 1 | 9 | |
| group 2 | 9 | |
| ***11. AO bridge cells*** | | **15** |
| ***12. Lamellate bodies*** | | **15** |
| group 1 | 4 | |
| group 2 | 4 | |
| group 3 | 6 | |
| group 4 | 1 | |
| ***13. Secretory cells C*** | | **13** |
| ***14. Lithocytes*** | | **8** |
| ***15. Secretory cells B*** | | **6** |
| ***16. Secretory cells A*** | | **4** |
| ***17. Large granular cells*** | | **4** |
| ***18. SNN*** | | **4** |
| **Total** | | **894** |

**Figure S2.** Extended table of the aboral organ cell types and cell numbers identified in the “M. leidyi RT” dataset.

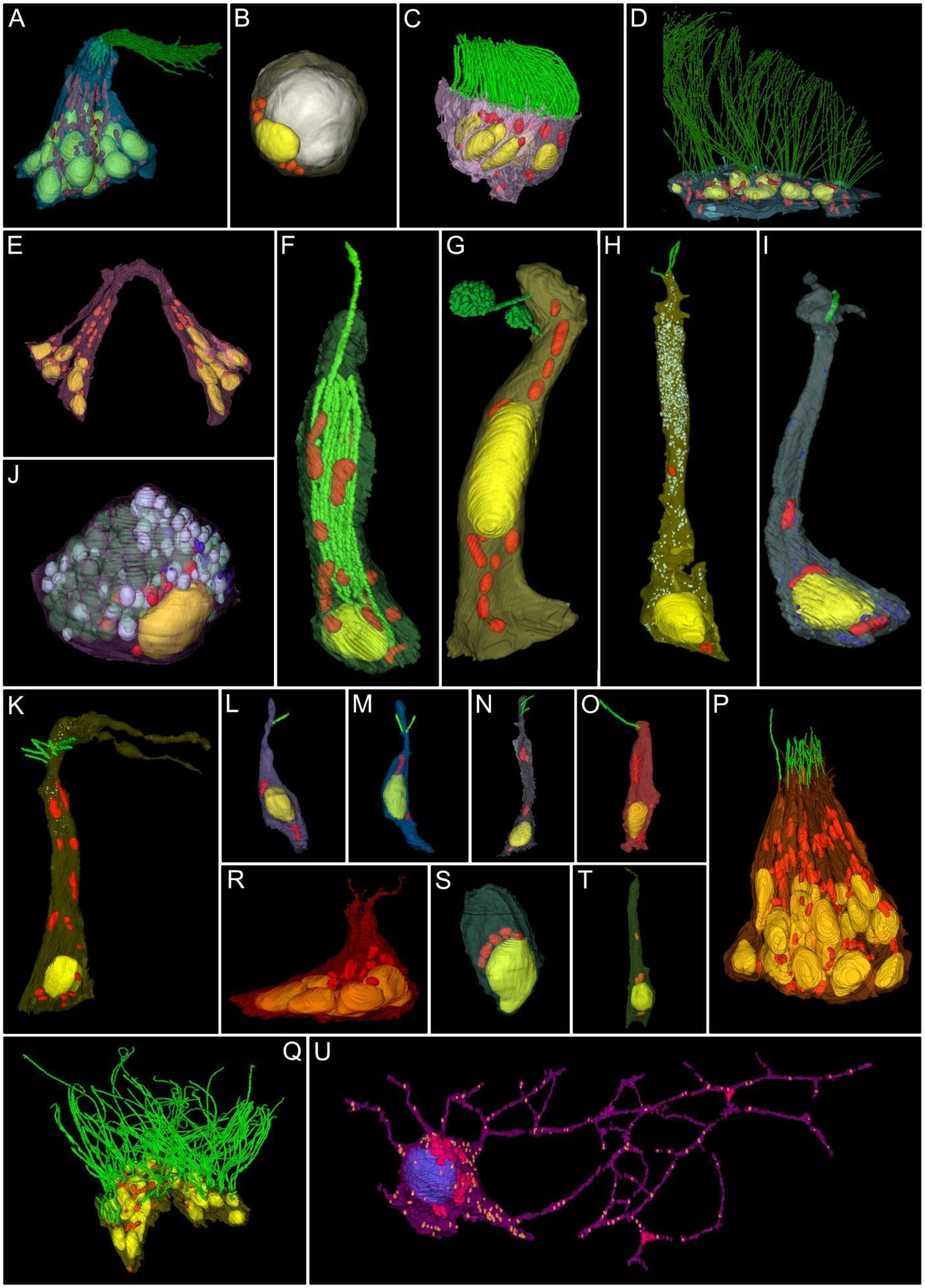
A
B
C
D
E
F
G
H
I
J
K
L
M
N
O
P
R
S
T
Q
U

**Figure S3. Cell type diversity in the *Mnemiopsis leidyi* aboral organ.** (A) Balancers. (B) Lithocyte. (C) Ciliated grooves. (D) Dome. (E) Bridge. (F) Lamellate body. (G) Pressure cell. (H) Putative secretory cells A. (I) Putative secretory cells B. (J) Large granular cells (large reflective granules are in pale green). (K) Putative secretory cells C. (L) Mono-ciliated cell. (M) Bi-ciliated cell. (N) Tri-ciliated cell. (O) Cell with longer, motile cilium. (P) Bundle cells. (Q) Polar fields. (R) Polar fields-associated cells. (S) Epithelial cells. (T) Epithelial elongated cell. Yellow: nuclei, red: mitochondria, green: cilia, light blue: electron-lucent vesicles, dark blue: electron-dense vesicles. (U) Neural cell body and neurites. Blue: nucleus; red: mitochondria; yellow: electron-dense vesicles.

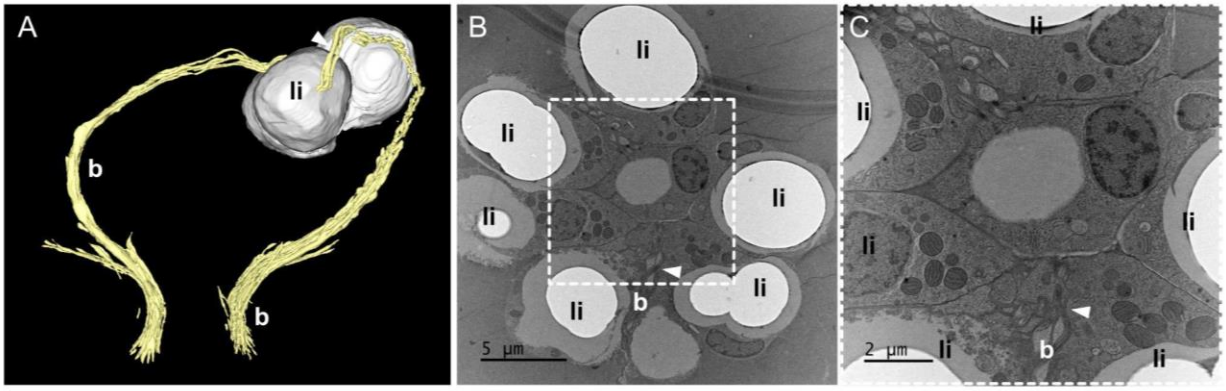


**Figure S4. Coupling of lithocytes with balancer cilia.** (A) 3D reconstruction of 2 groups of balancer cilia (pale yellow) and their interaction with two lithocytes (gray). The apical ends of the balancer cilia are tightly connected with the lithocytes (white arrowhead). (B) TEM micrograph of a lithocyte group and balancer cilia between two cells (white arrowhead). (C) Higher magnification TEM micrograph corresponding to the boxed area in (B). The white arrowhead points to the balancer cilia at the intersection of two lithocytes. (A to C) b: balancer cilia; li: lithocytes.

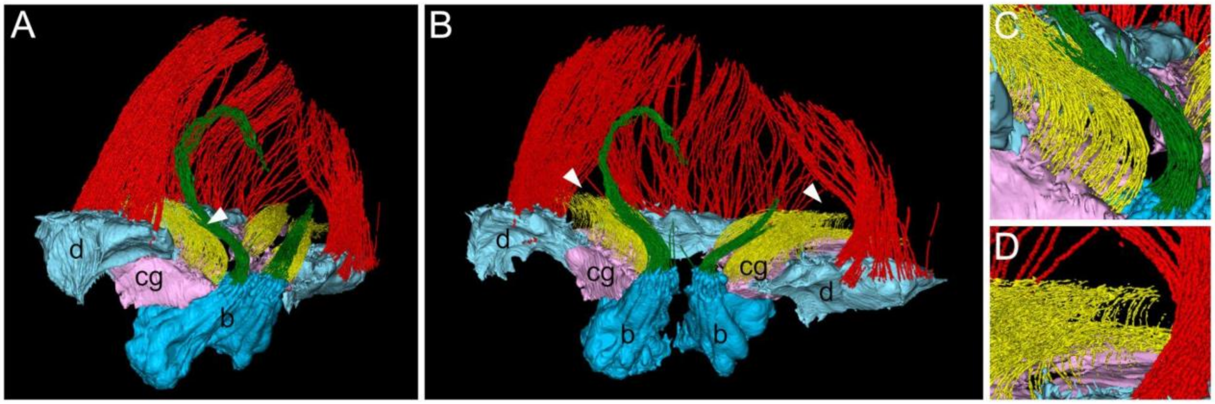


**Figure S5. Interplay of balancers, ciliated groove, and dome cilia.** (A) 3D reconstruction of balancers, ciliated grooves, and dome cells (lateral view). The apical sides of the ciliated groove cilia (yellow) are in contact with the basal side of the balancer cilia (green, white arrowhead). (B) 3D reconstruction as in (A). The dome cilia (red) form a “passage” (white arrowheads) which allows the ciliated groove to protract across the dome and extend towards the comb rows (not shown). (C) Details of the contact between the balancers and ciliated groove cilia. (D) Details of the ciliated groove protruding through the dome cilia. b: balancers; cg: ciliated groove; d: dome.

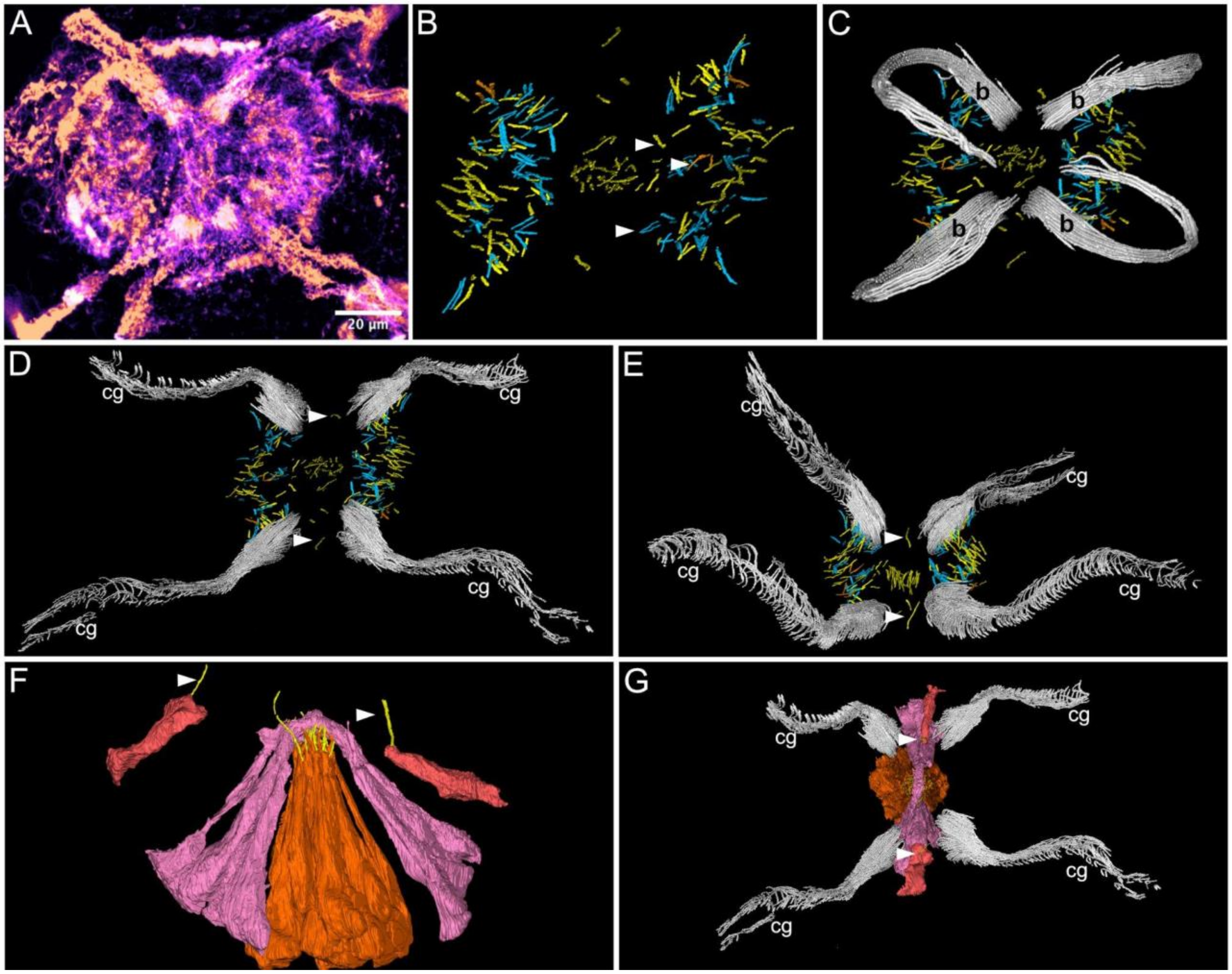


**Figure S6. Ciliated cells of the aboral organ.** (A) Maximum intensity confocal projection of sections through the aboral organ. The tubulin staining reveals the distribution of the cilia belonging to several ciliated cell types. (B) 3D reconstructions of SBFSEM data reveal the ciliary pattern of the aboral organ. The cilia are color-coded according to the cell types. The white arrowheads point to cilia belonging to mono-ciliated cells (in yellow), bi-ciliated cells (in blue), and tri-ciliated cells (in orange). (C) 3D reconstruction of the ciliary pattern as in (B), shown in combination with the balancer cilia (b, in gray). (D) 3D reconstruction of the ciliary pattern as in (B) along with the ciliated groove cilia (cg, in gray). The white arrowhead indicates a single motile cilium that can contact the adjacent ciliated grooves (cg). (A to D) top view. (E) Lateral view of the 3D reconstruction is shown in (D). The white arrowhead points to the motile cilium, which can interact with the adjacent ciliated groove cilia (cg). (F) 3D reconstruction of the cell bodies bearing the motile cilia (dark pink, white arrowheads), the bridge (pink), and the bundle cells (orange, lateral view). (G) 3D reconstruction showing the orientation of the bridge (pink), bundle cells (orange), and mono-ciliated cells (dark pink) with respect to the ciliated grooves (cg). The white arrowheads indicate the motile cilia.

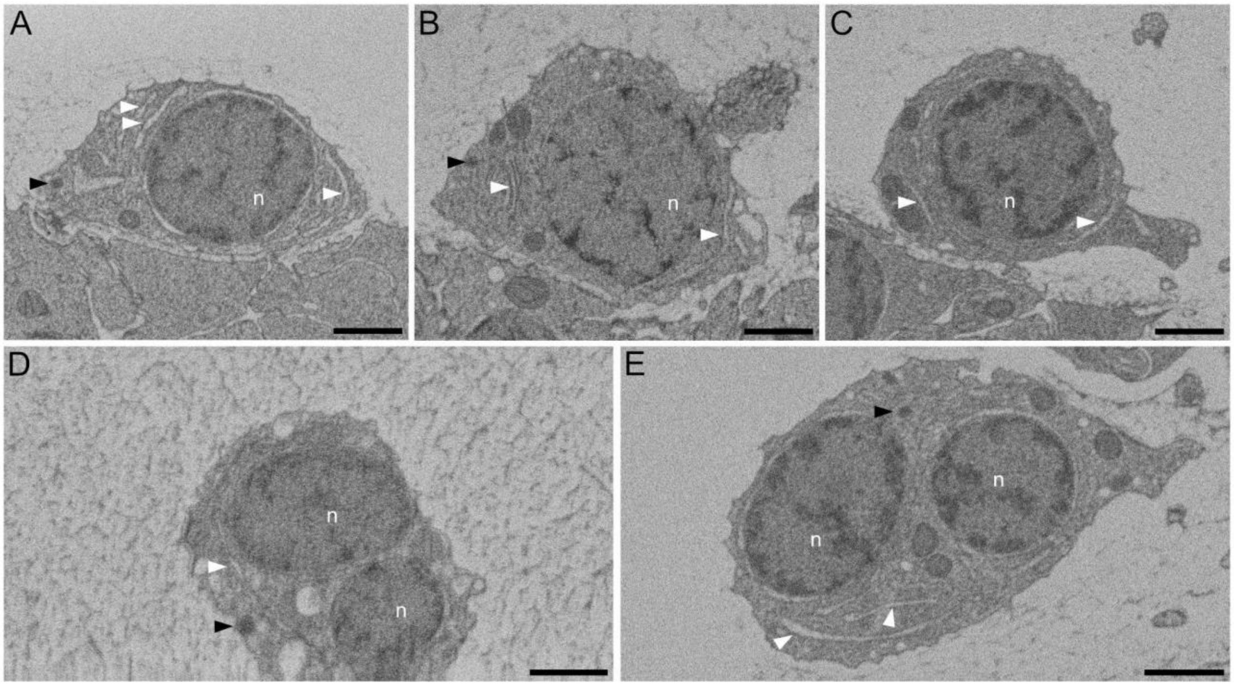


**Figure S7. Neuronal cell bodies of the aboral organ nerve net.** (A-E) SBFSEM micrographs of neuronal cell bodies from “M. leidyi HPF 2, 3, 4” datasets. (A-C) SBFSEM cross sections of neuronal cell bodies reveal several bundles of endoplasmic reticulum (white arrowheads) in the cytoplasm as well as dense core vesicles (black arrowheads). (D, E) SBFSEM sections of neuronal cell bodies, including two nuclei, bundles of endoplasmic reticulum (white arrowheads), and dense-core vesicles (black arrowheads). n: nuclei. Scale bars represent 1μm.

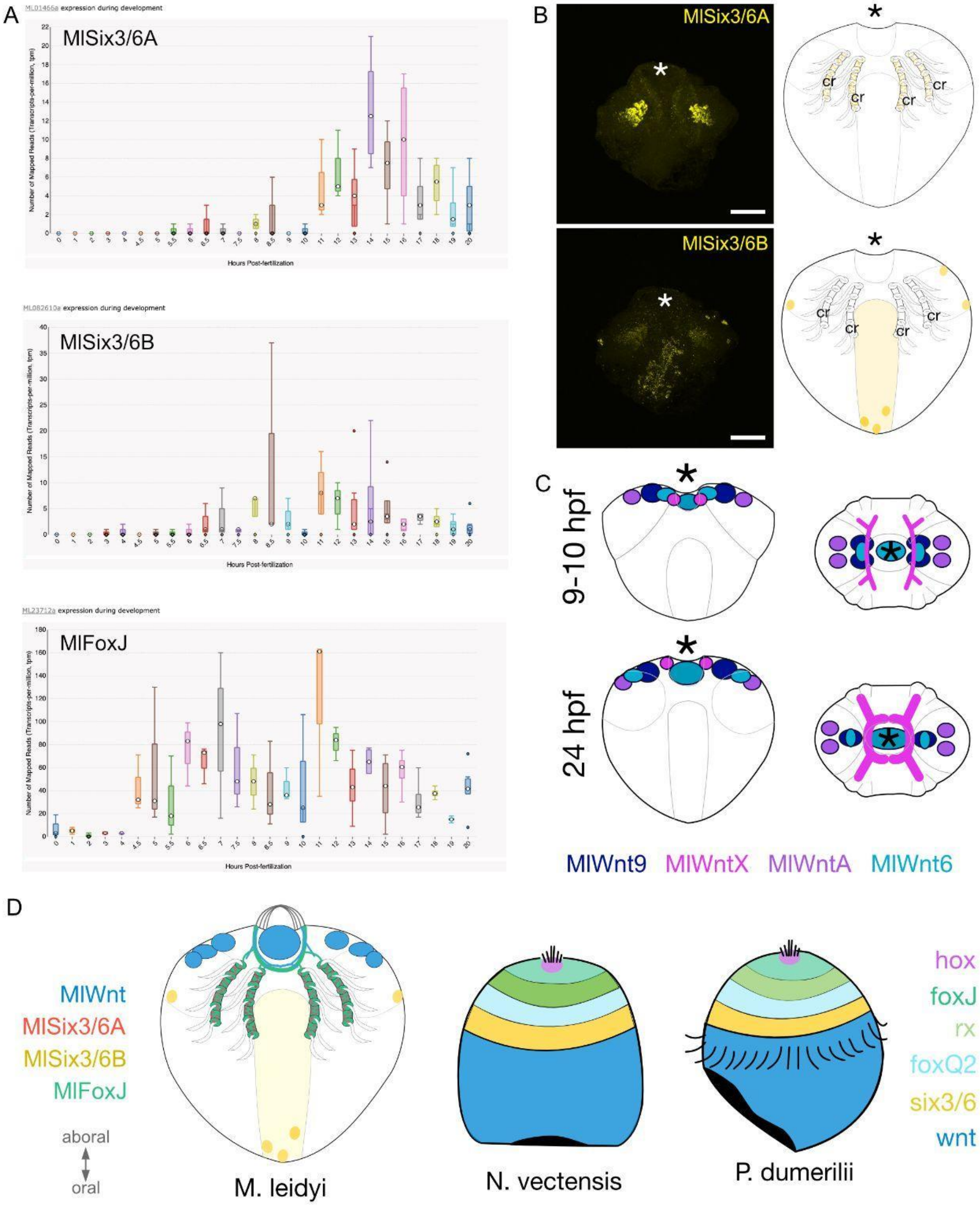


**Figure S8. Anterior-posterior patterning (AP) genes in *M. leidyi.*** (A) Temporal expression of candidates from 0 to 20 hpf. Expression plots are taken from https://research.nhgri.nih.gov/mnemiopsis/. (B) Expression patterns of one-day-old *M. leidyi* of *MLSix3/6A, MLSix3/6B,* and *MLFoxQ2* and diagrams. *MlSix3/6A* and *MlFoxQ2* were imaged using a 546 detector, whereas *MlSix3/6B* was imaged using a 647 detector. The signals appearing in the tentacle bulbs are non-specific. Asterisks indicate the position of the aboral organ. (C) Schematics of *MLWnt* gene expression from Pang et al, 2010. (D) Summary schematics including expression patterns of AP genes in *M. leidyi* (Pang et al., 2010 and this study)*, Nematostella vectensis,* and *Platynereis dumerilii.* Diagrams of *N. vectensis* and *P. dumerilii* summarize expression data from (12,37,39,67) and are adapted from Marlow et al., 2014.

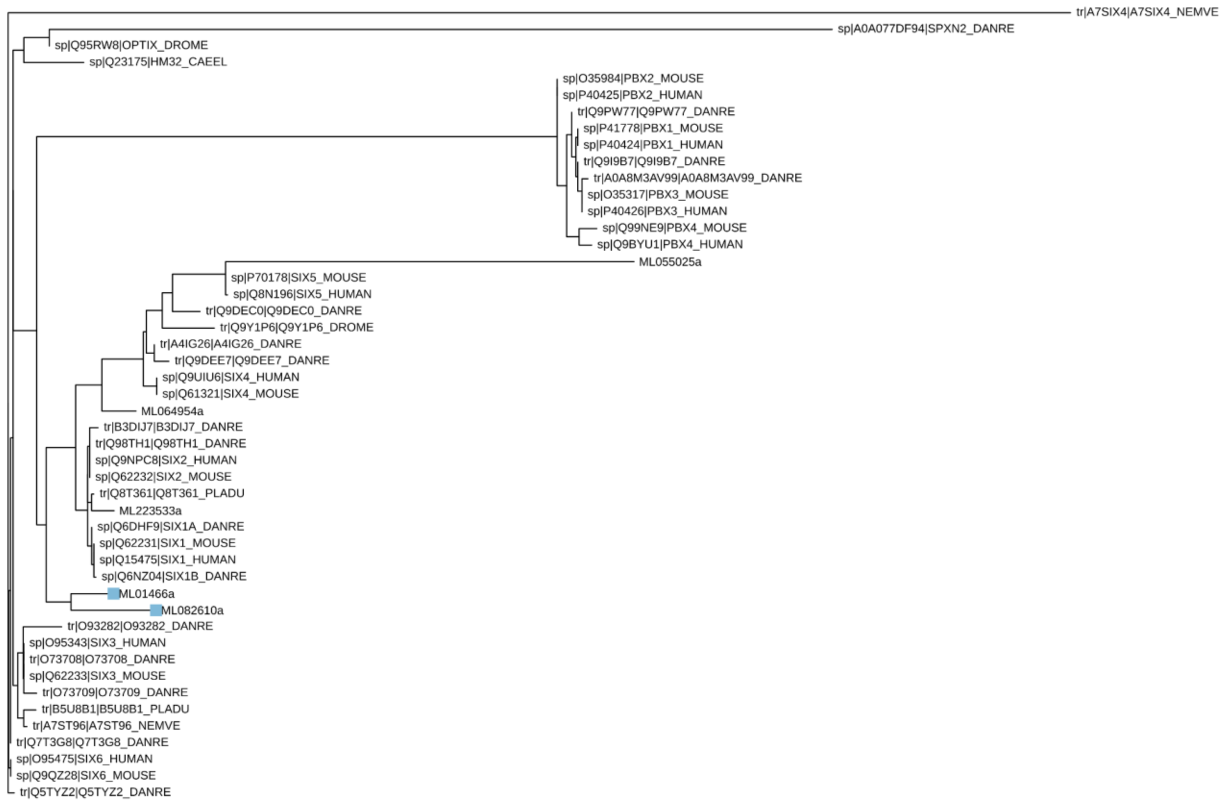


**Figure S9. Phylogenetic tree including *MLSix3/6A* and *MLSix3/6B***

*MLSix3/6A* and *MLSix3/6B* are highlighted in light blue.

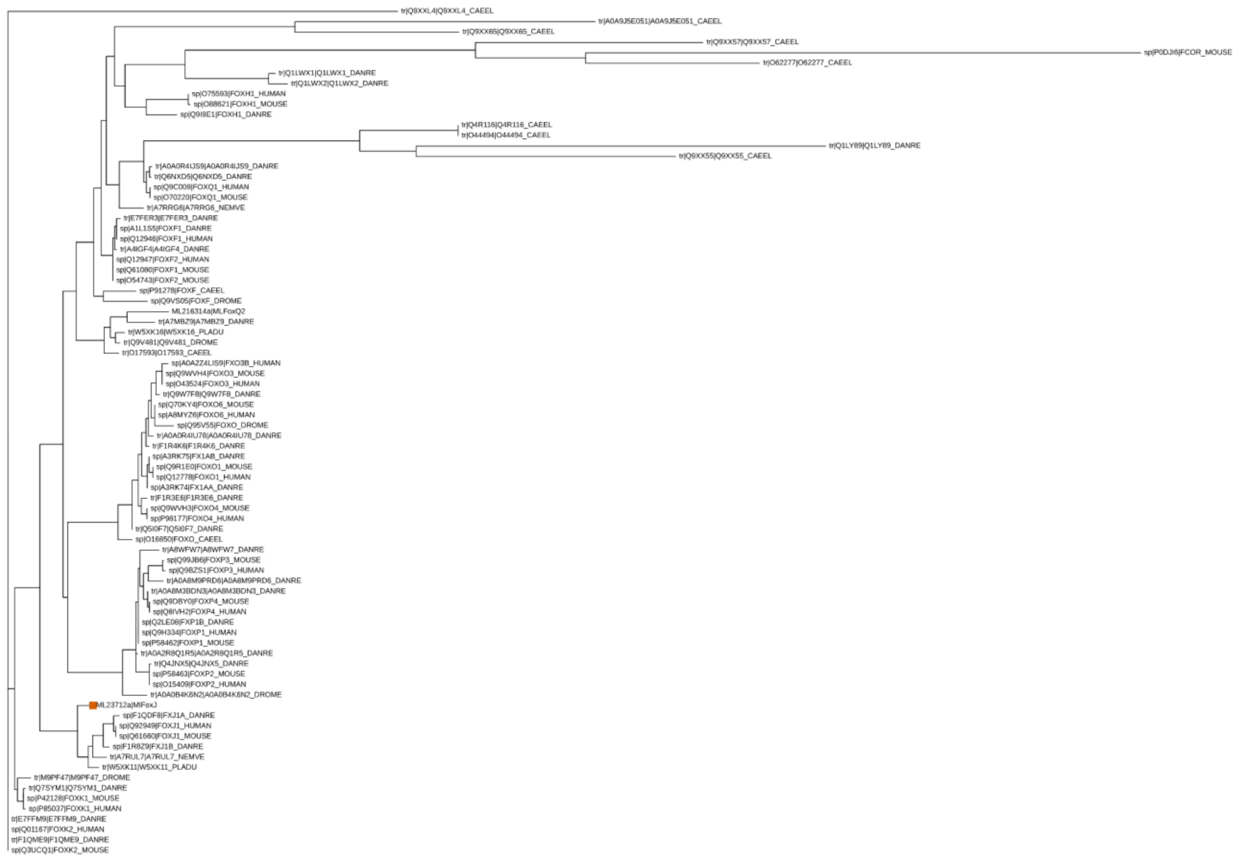


**Figure S10. Phylogenetic tree including *MLFoxJ***

*MLFoxJ* is highlighted in orange.
